# Malaria-derived hemozoin alters chromatin remodelling and skews dendritic cell responses to subsequent bacterial infections

**DOI:** 10.1101/2024.03.18.585548

**Authors:** Gintare Lasaviciute, Kanwal Tariq, Anaswara Sugathan, Jaclyn Quin, Ioana Bujila, Oleksii Skorokhod, Marita Troye-Blomberg, Eva Sverremark-Ekström, Ann-Kristin Östlund Farrants

## Abstract

Hemozoin (HZ), the malaria pigment, is released together with the with the parasite merozoites during the blood stage of the disease, which is connected with the pro-inflammatory response to induce T-cells and B-cells. Co-infections with bacteria lead to a more severe disease progression and the underlying mechanisms are poorly understood. Here, we investigated the impact of HZ on the early response of monocyte-derived dendritic cells (moDC) to a common bacterial component, LPS. A short-term HZ exposure for two hours did not induce an inflammatory response, but it did alter the transcriptional response to LPS. In moDC co-exposed to HZ and LPS, the induction of HLA-DR and PD-L1 gene expression was reduced and associated with decreased binding of RELA compared to LPS-stimulated cells. These gene promoters recruited the silencing chromatin remodelling complex NuRD upon co-exposure, instead of the PBAF complex at the promoter in LPS-stimulated cells. Further, HZ maintained transcription of C-type lectin receptors associated to an immature DC phenotype, DC-SIGN (CD209) and macrophage mannose receptor (MMR/CD206). Here, activated RELA and IRF3 were recruited in a PBAF and ncBAF dependent manner. Upon LPS co-exposure, NuRD was replacing these complexes to allow for a reduced transcriptional level of these immature markers. The association of chromatin remodelling complexes did not alter the chromatin state at the promoters, which was changed during differentiation to DCs or even at an earlier point. In conclusion, HZ exposure primes specific gene promoters at an early time point, which results in a different transcriptional response and may also lead to a changed immune reaction to bacterial co-infections.

**Graphical abstract:** 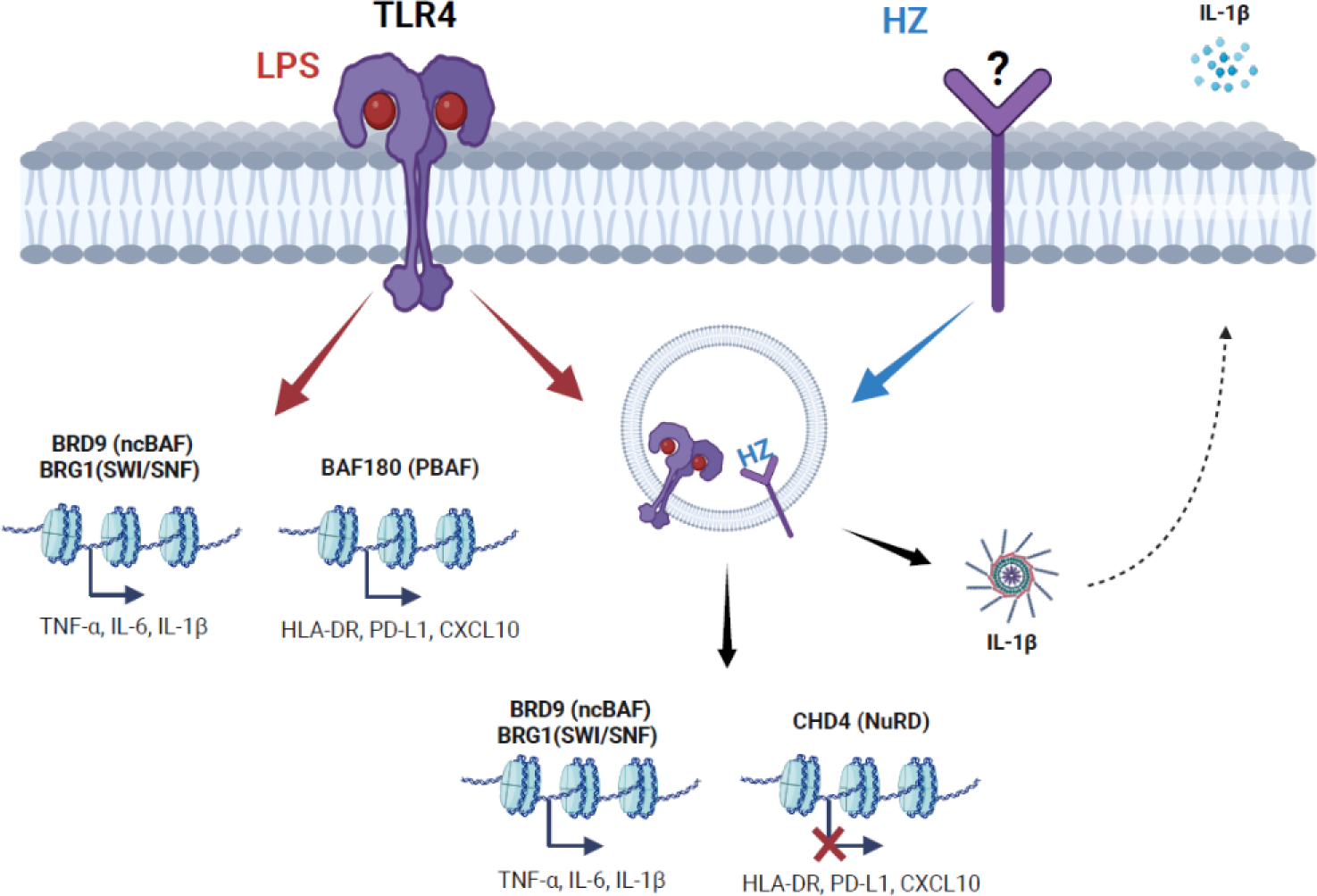

## Introduction

The human immune response to malaria is multifaceted and complex since the malaria parasite, Plasmodium, has many life stages and infect several organs in man. During the blood stage, infected erythrocyte releases the parasite merozoites together with erythrocyte and malaria specific content that elicit a strong innate pro-inflammatory response, which leads to activation of T-cells and to the clearance of the parasite (reviewed in 1). Co-infections with viruses or bacteria usually accelerate both the malaria and the infection, in particular in children and naïve infected individuals (2, 3).

The erythrocyte release contains several Plasmodium and host molecules (reviewed in 3 - 5), which all contribute to the immune response to malaria and maybe to the more sever disease progression in co-infections. Isolated macrophages or dendritic cells (DC) incubated with infected red blood cell lysate (iRBC) or purified hemozoin (HZ), a crystalline pigment from Plasmodium digested haem, do not elicit a pro-inflammatory response (6, 7) unless at high concentration (8). Other stimuli are also required and isolated cells require external ROS to mount a pro-inflammatory response (9, 10). Nevertheless, iRBC or HZ affects innate cells in other ways, as iRBC and HZ impairs DC function at several stages; the differentiation of monocytes to DC (moDC) *in vitro* (11, 12), moDC maturation upon exposure to stimuli (8, 11–15), and impaired T-cell activation (8, 13–16). However, the mechanism behind the interplay between pathways is not known.

Pre-incubation of DCs with DC for 24 hours or more affects subsequent responses to LPS, with an impaired maturation (8, 11). These effects may be a result of priming of innate cells to establish a trained or tolerance memory by epigenetic (histone modifications) and metabolic long-term reprogramming (18, 19). iRBC and HZ exposure induce a trained phenotype in monocytes from naïve blood donors, with a higher level of H3K4me3 at immune genes (19, 20). These changes work long-term by priming innate cells to subsequent infections, however, whether short-term exposure with HZ also has modulatory effects and alters the response to co-infections is poorly understood. iRBC have been shown to induce early responses in the acute phase of blood-derived DC, including secretion of chemokines but more prominently by altering the transcription of genes within a few hours (7). An early effect on transcription is observed in HZ exposed monocyte derived DC (moDC), and the early exposure time inhibited LPS induced maturation (14). The receptor for HZ is not identified or which pathways are activated. Nevertheless, when phagocytosed, HZ activates NFκB signalling in a MYD88 independent way (21–23).

Here, we show that exposure to HZ modulated the acute transcriptional response to LPS; co-exposure with HZ specifically prevented the upregulation by LPS of HLA-DR and PD-L1 gene expression, in contrast to inflammatory genes, where HZ had no effect on the LPS-induced activation. Many immune genes are regulated by NFκB and IRFs in an SWI/SNF dependent manner with specific constellations operating on different gene promoters (24, 25). We show that the HLA-DR and PD-L1 associated with the BAF180-PBAF variant upon LPS stimulation, and this complex was replaced by the silencing NuRD complex in co-exposed cells. NuRD antagonise the SWI/SNF on immune genes (26). Despite the association of chromatin remodelling factors, the regulation of these genes was not dependent on chromatin alterations. The genes in the DC response either established an open configuration during differentiation from monocytes to moDC, or was already pre-set in a very open state at an even earlier stage. Furthermore, HZ kept moDC in an immature state with maintained transcription of DC-SIGN and MMR genes (27). We propose that HZ primes specific gene promoters to respond differently to subsequent or simultaneous stimulation by bacteria through the recruitment of SWI/SNF complexes and NuRD, with a capacity to transiently alter a following T-cell response. At the same time transcription is actively maintained on immaturity genes important for homeostasis.

## Material and Methods

### Human subjects

Peripheral blood cells were retrieved from anonymous, healthy blood donors who cannot be traced back; thus, the project does not require approval from the Swedish ethical review authority.

### Natural Hz preparation

Natural HZ was prepared as previously described (14, 15). Briefly, 12 h after schizogony of synchronized FCR3 parasite cultures, the supernatant was collected. A 10%/40% interphase of a 6% mannitol-containing Percoll gradient was used to collect natural HZ, which was then extensively washed with 10 mM phosphate buffer followed by washing with PBS. HZ was opsonized with equal volume of human AB serum for 30 min at 37°C and then it was quantified according to its haem content. HZ preparation with 4 nmol/μl haem content was used in the study. Prior to use, HZ was resuspended by passage through a 0.4 mm needle fitted to a 1 ml syringe.

### Cell culture and stimulations

Peripheral blood mononuclear cells (PBMC) were isolated from buffy coats using Ficoll-Hypaque (Cytiva) gradient centrifugation. Monocytes were enriched from PBMC by negative selection using EasySep^TM^ human monocyte kit (STEMCELL Technologies) according to the manufactureŕs instructions. Monocytes were resuspended at 1×10^6^ cells/ml and were seeded in 96-well plates (with 200 μl/well) or 6-well plates (with 2 ml/well) using RPMI-1640 culture medium which was supplemented with 20 mM HEPES, 2 mM L-glutamine, 100 U/ml penicillin, 100 μg/ml streptomycin (all from Cytiva), 5% human AB serum (Sigma-Aldrich), 50 μM 2-mercaptoethanol, 2% sodium pyruvate (both from Gibco), 35 ng/ml IL-4 and 50 ng/ml GM-CSF (both from PeproTech). On day 3, half of the medium volume was replaced with the fresh medium containing IL-4 and GM-CSF. On day 5, cells were pre-exposed to HZ (5 μl for 96-well plates and 50 μl for 6-well plates) and the plates were centrifuged at 700 r.p.m. for 20 s in order to initiate the contact with Hz. After 2 h, 50 ng/ml of LPS from Escherichia coli O26:B6 (Sigma-Aldrich) was added to appropriate wells for additional 2 h or 24 h. Cells assigned to HZ treatment alone were exposed to HZ at the same time as was added LPS. Cells kept in complete culture medium served as the control. Supernatants were collected and stored at −20°C until further analysis, while cells were collected either for western blot, chromatin immunoprecipitation (ChIP), mRNA expression analysis, ATAC-qPCR or flow cytometry. Throughout the experiment, cells were incubated at 37°C with 5% CO_2_.

### Immunoblotting

MoDC were harvested in 4xLameli buffer, separated on a 12% PAGE and transferred to a PVDF (Miilipore) membrane. The membrane was blocked in 5% milk and incubated separately with primary antibodies: anti-Phospho-IRF3 (Ser 396) (D601M, Cell Signaling), anti-IRF3 (D6I4C, Cell Signaling), anti-Lamin B1 (ab65986, Abcam), anti-Caspase-1 (ab207802, Abcam), anti-beta Tubulin (ab6046, Abcam). Immunoblots were developed using ECL Western Blot Substrate (BioRad). The quantification of protein signals was performed by ImageLab^TM^ (ChemiDoc^TM^ Imaging system, BioRad) software by comparing the signals from each target proteins to control proteins.

### Chromatin immunoprecipitation

Chromatin from different treatment groups was cross-linked using 1% formaldehyde (Sigma-Aldrich). Chromatin from 1×10^6^ cells was incubated with antibodies against histone H3 (ab1791), histone H3K27Ac (ab4729), histone H3K4me3 (ab8580), RelA (NFκB p65, ab7970), IRF3 (ab76409) IgG (ab171870) (all from Abcam), or BRG1 antibody (generated as previously described, 28). Protein A/G magnetic beads (ThermoScientific^TM^) were used to precipitate. The target DNA was amplified by qPCR using KAPA SYBR FAST qPCR Master mix (Kappa Biosystems Inc). The results are presented as percentage of input, with the IgG control subtracted from the Ct values obtained for samples. Primers are listed in the Table S1.

### RNA extraction and real-time quantitative polymerase chain reaction

Total RNA was extracted using TRI reagent (ThermoScientific^TM^), and reverse transcription of RNA was performed using SuperScript VILO cDNA synthesis kit (ThermoScientific^TM^), according to the manufacturerś instructions. qPCR was performed using KAPA SYBR FAST qPCR Master mix. mRNA expression levels were calculated using 2^-ΔΔCt^ method and PP1A was used for normalization. Primer pair list can be found in the Table S1.

### ATAC-qPCR

Cells were collected and tagged with Tn5 transposase (Illumina) according to the manufacturer’s instructions. DNA was purified using MinElute PCR purification kit (Qiagen) and it was stored at −20°C until further analysis by qPCR. Tagmentation-specific forward primer and gene-specific reverse primer was used for amplification. IL-2 was used for normalization. Primer pair list can be found in the Table S1.

### Flow cytometry

Following cell collection on day 5 or day 6, moDC were stained with alternating combination of antibodies: CD11c APC (clone 3.9), MMR PerCP-Cy5.5 (clone 15-2), TLR4 PE (clone HTA125), CLEC12A FITC (clone 50C1), CD83 PeCy-7 (clone HB15e), HLA-DR PerCP (clone L243), PD-L1 PE (clone 29E.2A3) (all from BioLegend), CD14 PeCy7 (clone MφP9), DC SIGN BV421 (clone DCN46) and CD86 FITC (clone FUN-1) (all from BD Biosciences). A fixable viability stain 780 (BD Biosciences) was used to exclude non-viable cells. Samples were acquired using FACSVerse flow cytometer (BD Biosciences) and FlowJo Software (TreeStar) was used to analyse the data.

### Cytometric bead array (CBA)

Soluble levels of TNF-α, IL-10, IL-6 and IL-1β cytokines were measured in the cell culture supernatant after 2 h of exposure using CBA human Inflammatory cytokine kit (BD Biosciences), while chemokines CXCL8/IL-8, CCL5/RANTES, CXCL9/MIG, CCL2/MCP-1 and CXCL10/IP-10 were quantified using CBA human chemokine kit (BD Biosciences), according to the instructions from the manufacturer. Samples were acquired using FACSVerse flow cytometer and FlowJo Software was used to analyse the data.

### Enzyme-linked immunosorbent assay

Soluble levels of TNF-α, IL-6, IL-1β, IL-23 and IL-10 (MabTech AB), as well as IL-RA (R&D systems, Bio-Techne), were measured in the cell culture supernatant after 2 h and/or 24 h of co-exposure according to the instructions from the manufacturer. The results were analysed using SoftMax Pro 5.2 rev C (Molecular Devices Corp.).

### Statistical analysis

Statistical analysis was carried out using GraphPad Prism 10 (GraphPad Prism Inc.). Non-parametric Friedman ANOVA test or non-parametric Kruskal-Wallis ANOVA test, followed by Dunńs Multiple Comparison Test was applied to determine the differences between multiple treatment conditions. Non-parametric Wilcoxon matched-pairs signed rank test or non-parametric Mann-Whitney test was used to determine the differences between two treatment conditions. Statistical differences were considered significant if p-values were <0.05.

## Results

### HZ compromises LPS-induced upregulation of HLA-DR and PD-L1 gene expression

To address the question whether HZ has an impact on the early moDC phenotype and functional response to an unrelated microbial compound such as the bacterial-derived LPS, we pre-incubated cells with HZ for 2 h to allow its phagocytosis before adding LPS as depicted in Figure 1A. To mimic the state of HZ after its release in bloodstream, we used an opsonized preparation in which the HZ crystal contained plasma components. HZ was phagocytosed by the cells already after 2 h, as shown by the increased granulation of cells observed by flow cytometry (Supplementary Figure S1A). The exposure to HZ for 2 h did not affect the surface expression of HLA-DR and it did not hamper LPS-induced expression of this marker after 24 h (Figure 1B). As expected, the expression of the maturation marker CD83 was significantly upregulated by LPS after 2 h and 24 h (Figure 1B). Since HZ mainly impacts the early transcriptional response (14), we investigated mRNA levels of these DC markers upon LPS-stimulation. Interestingly, the LPS-triggered increase in HLA-DR transcripts was markedly reduced when the cells was co-exposed to HZ (Figure 1C). Transcription of CD83, however, was not reduced by co-exposure of HZ and LPS compared to LPS alone. This was even more pronounced for the gene expression of the maturity marker CCR7, which was induced by LPS and further increased when the cells were co-exposed to HZ (Figure 1C).

**Figure 1.**
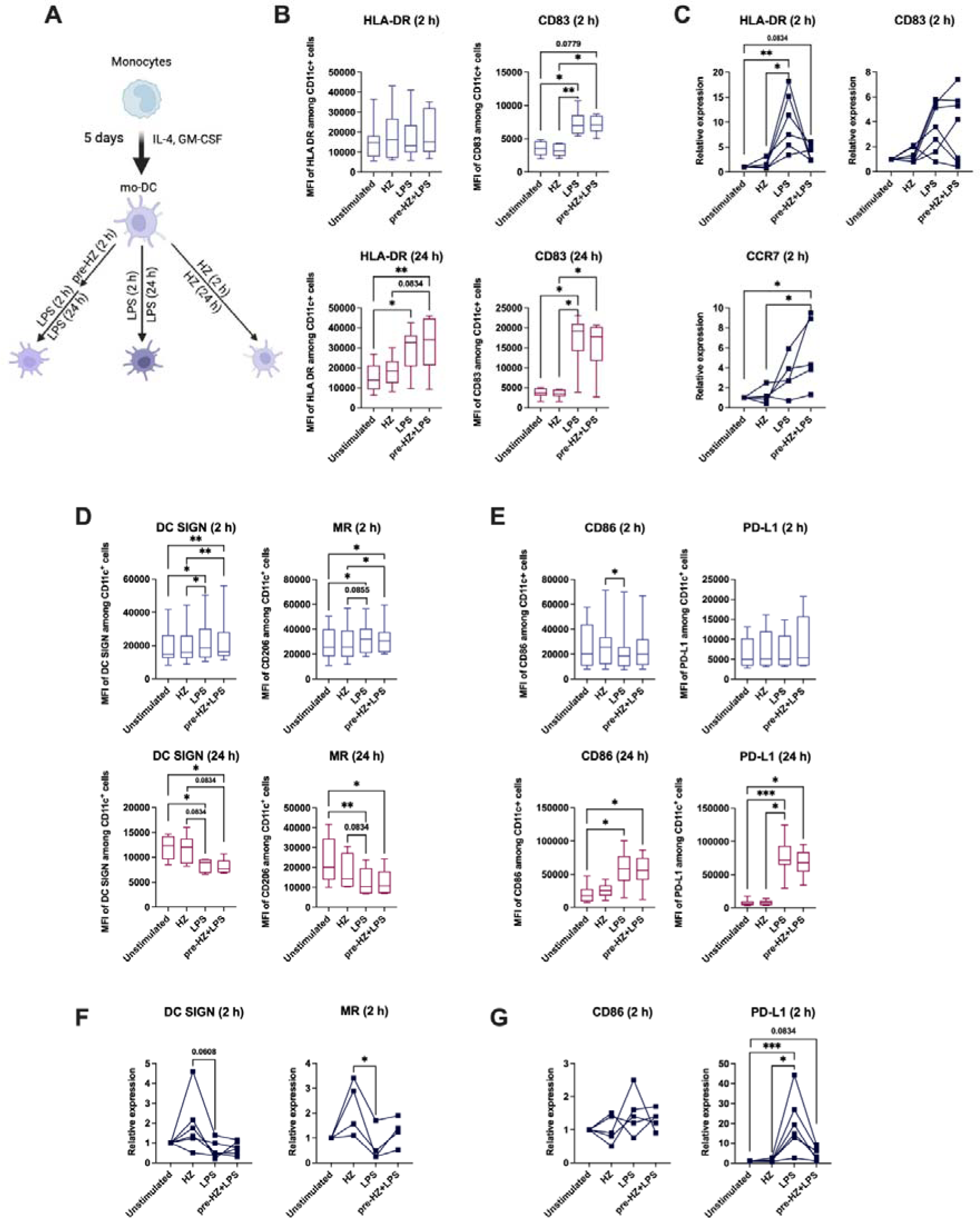
HZ prevents the upregulation of HLA-DR and PD-L1 transcripts during co-exposure with LPS. **(A)** Experimental model of monocytes differentiation to mo-DC during 5 days culture and exposure to different stimuli. MoDC were pre-exposed to HZ for 2 h, then LPS was added for additional 2 h or 24 h (co-exposure condition). Cells kept in complete culture medium served as the control. **(B)** The mean fluorescent intensity (MFI) of HLA-DR and CD83 in moDC after 2 h exposure (upper graphs) and 24 h exposure (lower graphs). **(C)** The relative mRNA expression of HLA-DR, CD83 and CCR7 in moDCs after short-term exposure. The mean fluorescent intensity (MFI) **(D)** DC SIGN and MMR, **(C)** CD86 and PD-L1, expression in moDC after 2 h exposure (upper graphs) and 24 h exposure (lower graphs). The relative mRNA expression of **(F)** DC SIGN and MMR, **(G)** CD86 and PD-L1, in moDC after short-term exposure. (B-G) Paired Friedman test followed by Dunńs multiple comparison was used to determine statistical difference, n.s.=p>0.05, *p<0.05, **p<0.01, ***p<0.001, n=4-12.

To further assess if moDC remained immature with poor T-cell activation capacity upon HZ addition, we investigated the surface expression of the C-type lectin receptors DC-SIGN (CD209) and MMR (CD206), which are associated with an immature phenotype (27). The surface expression of both receptors in HZ exposed cells was high and comparable to the unstimulated cells, while LPS caused a downregulation of these receptors after 24h, which was not prevented by HZ pre-exposure (Figure 1D). Further, HZ did not block the LPS-mediated upregulation of co-stimulatory receptor CD86 or the surface expression of PD-L1, which is a negative co-stimulatory receptor important for T-cell inhibition, following 24 h (Figure 1E). To assess the early transcriptional level of these factors, we examined the mRNA levels of DC SIGN and MMR. The mRNA levels were reduced by LPS compared to the cells only exposed to HZ (Figure 1F). The mRNA levels of PD-L1 displayed a similar pattern to HLA-DR; co-exposure with HZ strongly reduced the expression of LPS-induced PD-L1 (Figure 1G). CD86 did not differ among samples at this early time point (Figure 1G). Taken together, our results suggest that HZ interferes with the stimulation by bacterial antigen during the early induction of gene expression specifically of genes important in antigen presentation and activation of T-cells.

### HZ does not influence the secretion of LPS-induced inflammatory and anti-inflammatory factors early during the response

We next investigated whether HZ had an impact on the LPS-induced inflammatory and anti-inflammatory protein production. As expected, LPS induced TNFα and IL-6 release already after 2 h, while IL-23 secretion was induced by LPS after 24 h, regardless whether HZ was present or not (Figure 2A and Supplementary Figure S2A). LPS also induced a strong transcriptional response of the TNFα, IL-6, and 1L-23 genes after 2 hours, which was not interfered with by HZ (Figure 2B and Supplementary S2A, lower panel). Furthermore, HZ did not interfere with the LPS-induced secretion of CXCL8, CXCL9 or CXCL10 chemokines (Figure 2C). CCL2 was not induced above background, while CCL5 was slightly upregulated in co-exposed cells (Supplementary Figure S2B). At the transcriptional level co-exposure resulted in enhanced CXCL9 expression, while CXCL10 resembled a similar pattern to that of HLA-DR and PD-L1; the increase was only detected in LPS exposed cells but not in LPS and HZ co-exposed cells (Figure 2D). For the other chemokines, mRNA expression varied substantially at 2 h (Figure 2D and Supplementary Figure S2B, lower panel).

**Figure 2.**
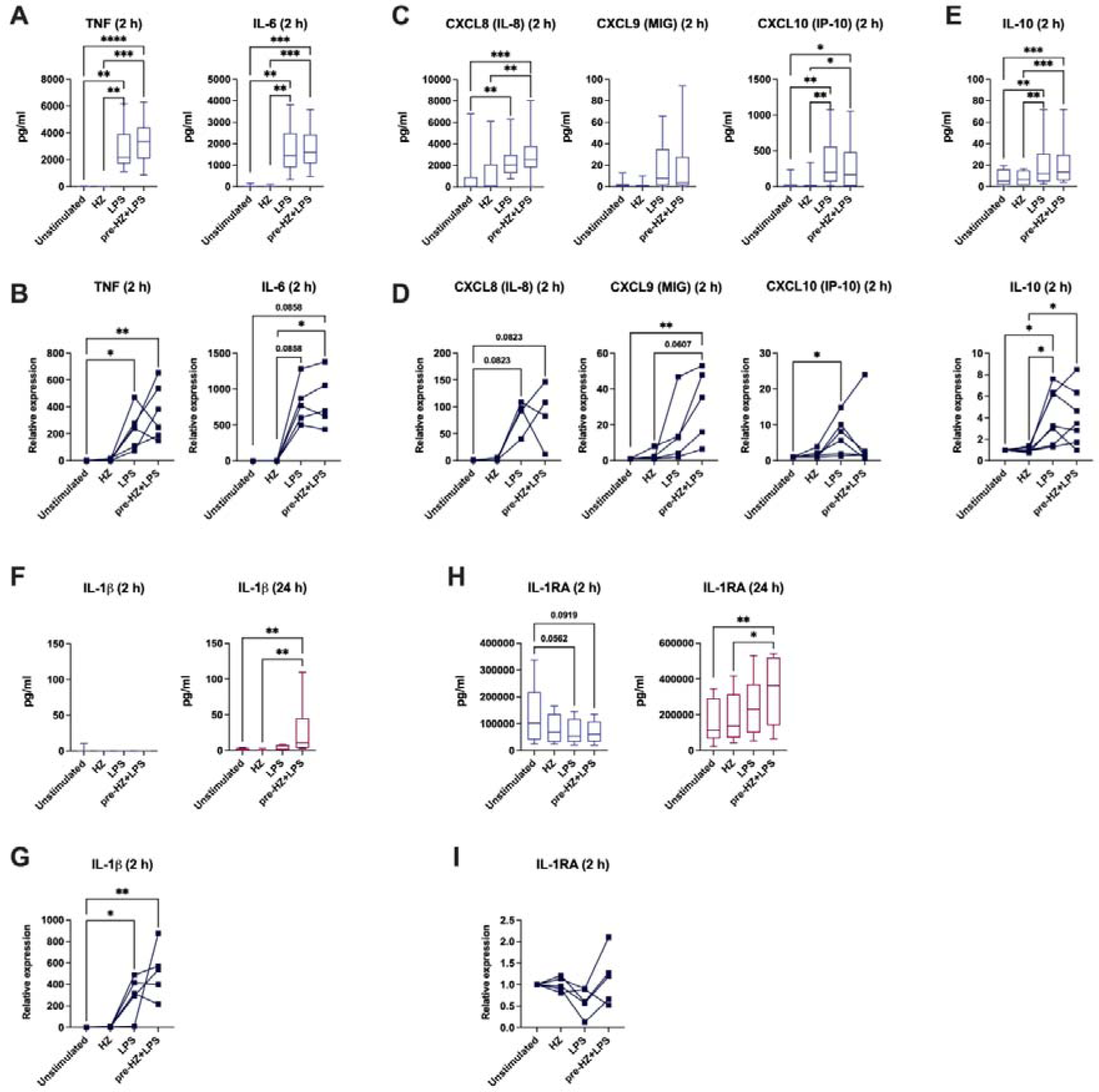
HZ does not interfere with the upregulation of inflammatory gene transcripts, but both HZ and LPS are required for the release of inflammasome-derived IL-1β. . **(A)** Soluble levels of TNF-α and IL-6 in moDC after 2 h exposure. **(B)** Relative mRNA expression of TNF-α and IL-6. **(C)** Soluble levels of CXCL8, CXCL9, CXCL10 after 2 h exposure. **(D)** Relative mRNA expression of CXCL8, CXCL9, CXCL10. **(E)** Soluble levels and relative mRNA expression of IL-10 after 2 h exposure. **(F, H)** Soluble levels of IL-1β and IL-1RA after 2 h and 24 h exposure. **(G, I)** Relative mRNA expression of IL-1β and IL-1RA. mRNA levels were measured in all samples after 2 h of exposure. (A-I) Paired Friedman test followed by Dunńs multiple comparison was used to determine statistical difference, n.s.=p>0.05, *p<0.05, **p<0.01, ***p<0.001, ****p<0.0001, n=4-14.

HZ did not interfere with the anti-inflammatory response as LPS-induced IL-10 production both at the protein and transcriptional levels 2 h post exposure (Figure 2E). Further, HZ did not have clear effects on the expression of the immunosuppressive or tolerance markers SOCS2 and IDO1, or reduced the LPS induction in co-exposed cells (Supplementary Figure S2C). We also investigated the protective enzyme Haem oxygenase (HO-1), as it might be induced upon haem release and dampen the response (29). HZ exposure did not trigger the gene expression of HO-1, regardless of LPS presence (Supplementary Figure S2C). In summary, HZ did not induce an inflammatory response or interfere with the response by LPS regarding secretion of cytokines and chemokines, nor with the transcriptional induction of the genes.

### HZ accounts for the release of inflammasome-derived IL-1β during co-exposure with LPS

Since HZ exposure has been associated with NLRP3-inflammasome activation (30, 31), we measured the inflammasome-dependent cytokine IL-1β secretion. The cytokine was detected only from cells co-exposed to HZ and LPS after 24 h (Figure 2F), preceded by enhanced levels of transcription by LPS stimulation as detected at 2 h (Figure 2G). In addition, transcriptional induction of the inflammasome component NLRP3 was upregulated by HZ and LPS co-exposure and by LPS stimulation alone, while the pro-caspase-I exhibited inductions by LPS alone (Supplementary Figure S2D). This suggests that the release of the IL-1β requires HZ to be co-exposed with a pathogen which induces a transcriptional response of inflammasome associated genes. The inflammasome activation involves the cleavage of pro-caspase 1 and we could not detect any cleavage as early as 2 h after stimulation, showing that 2 h is not sufficient for the full activation of the inflammasome to release IL-1β (Supplementary Figure S2E). During inflammation, the production of IL-1β is balanced by the IL-1 receptor antagonist (IL-1RA) (32). IL-1RA release was lower in LPS exposed and co-exposed cells after 2 h (Figure 2H), similar to IL-1RA mRNA expression at 2 h (Figure 2I). Still, regardless of HZ-exposure LPS-stimulation resulted in an increase of IL-1RA secretion after 24 h (Figure 2H). Our results indicate that HZ is responsible for the IL-1β inflammasome response during co-stimulation with LPS, and LPS responsible for the required induction of IL-1β and NLRP3 gene expression and the IL-1β secretion this is accompanied by a higher IL-1RA release.

### HZ does not interfere with the LPS activation by TLR4

To investigate the mechanism behind the dampening effect of HZ on the transcriptional response of HLA-DR and PD-L1 but not on inflammatory genes, we examined whether HZ changed the level of LPS activation through TLR4. The percentage of TLR4 expressing cells significantly decreased after 2 h both in cells stimulated with LPS and cells exposed to LPS and HZ (Figure 3A), showing that HZ did not affect receptor internalisation. The receptor level was restored at 24 hours, even significantly upregulated in co-exposed cells (Figure 3A). The receptor for HZ is still not known, but CLEC12A, the receptor for uric acid has been suggested to be the receptor for phagocytosis of HZ (33). The surface level of CLEC12A did not change in any treatments (Supplementary Figure S3A).

**Figure 3.**
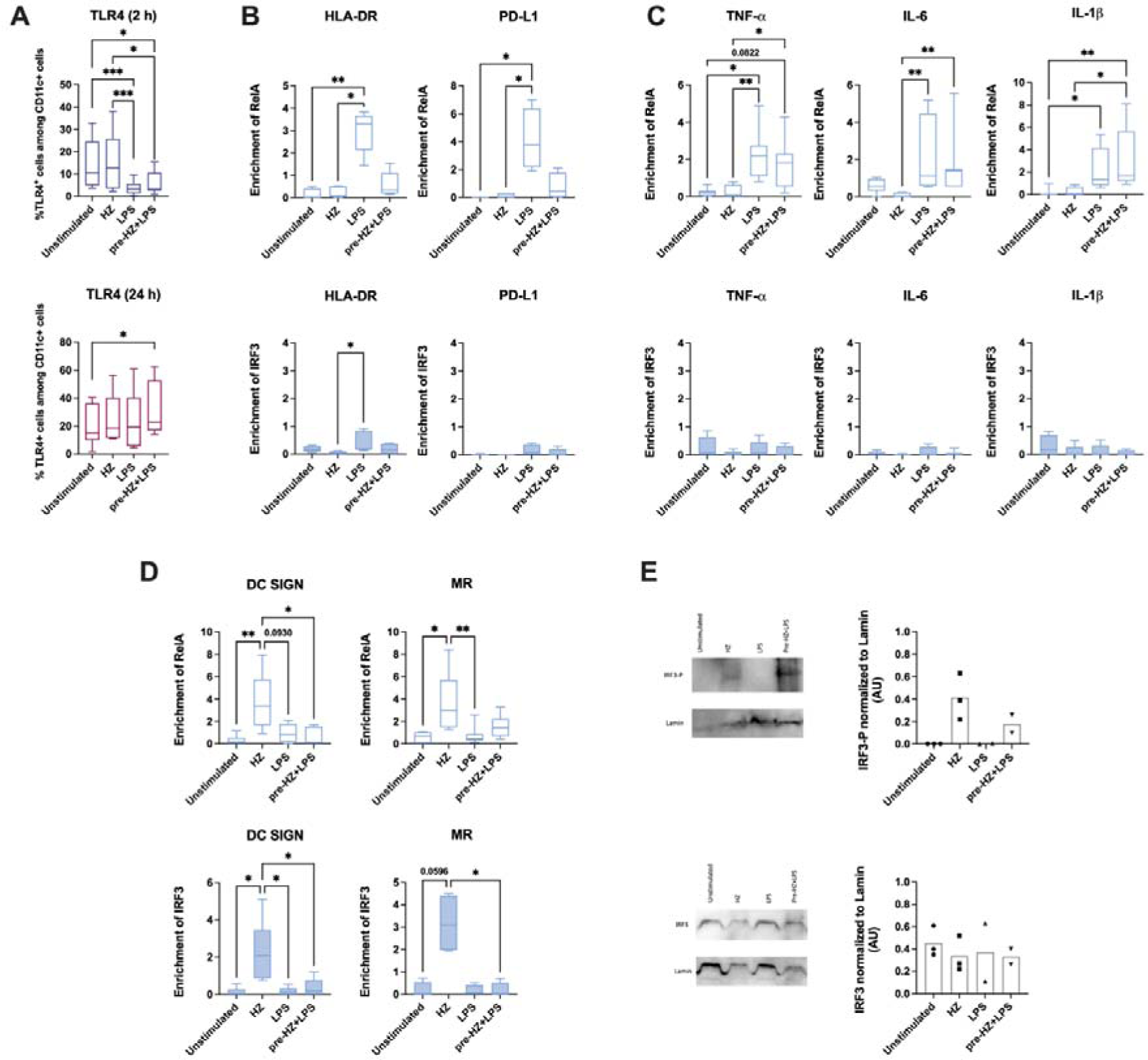
HZ influences transcription factor recruitment to the promoters of moDC genes after short term exposure. **(A)** The percentage of TLR4 expressing cells among CD11c^+^ moDC population. Paired Friedman test followed by Dunńs multiple comparison was used to determine statistical difference, *p<0.05, **p<0.01, ***p<0.001. The enrichment of RelA and IRF3 at the promoters of **(B)** HLA-DR and PD-L1, **(C)** TNF-α, IL-6 and IL-1β, **(D)** DC SIGN and MMR. (B-D) Non-parametric Kruskal-Wallis ANOVA test, followed by Dunńs multiple was comparison was applied to determine significant differences, n.s.=p>0.05,*p<0.05, **p<0.01. The data are present as median with interquartile range. N=4-12 **(E)** Representative immunoblot of the IRF3 protein and its phosphorylated form and quantitative analysis of each to the right. IRF3 levels were normalized to lamin levels (n=2-3).

### HZ phagocytosis changes the transcription factor recruitment to promoters

TLR4 ligation activates several transcription factors including the NFκB factor RELA and IRF3. To investigate the differentiated gene response to HZ in co-exposed cells, we examined the binding of these factors to the promoters of HLA-DR and PD-L1 genes. These promoters bound significant levels of RELA in cells stimulated with only LPS, while HZ and LPS co-exposed cells displayed markedly lower levels of binding (Figure 3B, upper panel). IRF3 was only recruited at low levels in all samples (Figure 3B, lower panel). In contrast, HZ did not hamper the binding of RELA at the promoters of inflammatory genes in LPS stimulated and HZ and LPS co-exposed cells, as it was significantly recruited to the promoters of the primary response gene TNFα and the secondary response genes IL-6 and IL-1β genes (Figure 3C, upper panels). A similar trend for RELA was also observed for IL-23 and CCL5 genes (Supplementary Figure S3B). IRF3 did only bind at low levels to the promoters of the inflammatory genes (Figure 3C, lower panels), even to the IRF3 target gene CCL5 after 2 hours (Supplementary Figure S3B). The RELA and IRF3 recruitment on the CD83 promoter resembled that of HLA-DR and PD-L1 with RELA occupancy in LPS stimulated cells, and low levels of IRF3, although no compromised transcription was observed in co-stimulated cells (Supplementary Figure S3C).

The slightly lower mRNA levels of DC-SIGN and MMR in LPS stimulated cells compared to HZ exposed cells (Figure 1F), prompted us to also assess transcription factor binding at these promoters. The difference in expression depended on transcription factor recruitment to DC-SIGN and MMR and both RELA and IRF3 were bound to the promoters of DC-SIGN and MMR in HZ stimulated cells (Figure 3D), a binding that was counteracted in LPS-stimulated cells (Figure 3D). To further examined the transcriptional effect on these genes, we explored published RNA seq data from DCs after 3h of exposure (7). This data clearly shows an induction of DC-SIGN by iRBC, whereas the transcript level in LPS-exposed cells remained at the unstimulated level. Taken together, this indicates that HZ not only impairs the recruitment of activating factor at certain genes, but also activates other groups of gene to maintain the level of transcription without hampering the regulation by other stimuli.

Since IRF3 was bound to the promoters of DC-SIGN and MMR in HZ exposed cells after 2 h, we examined the activation of IRF3 in the different treatments. IRF3 was activated, as shown with the presence of phosphorylated IRF3, in cells exposed to HZ alone and cells co-exposed with LPS and HZ (Figure 3E), showing that HZ exposure had an immune modulatory effect by activating both RELA and IRF3.

### Exposure to HZ does not alter chromatin states

The differential binding of RELA and IRF3 to gene promoters in exposed cells motivated us to investigate the chromatin states at the different gene promoters. No significant changes were detected in the chromatin accessibility analysed by ATAC-qPCR; HLA-DR and PD-L1 displayed no altered accessibility at the gene promoters or at gene enhancers by any exposure (Figure 4A). Nor did changes occur at the promoters, enhancers or gene bodies of inflammatory genes or lectin receptor genes, despite the large transcriptional induction in inflammatory genes (Supplementary Figures S4A and S4B). Many of these genes are also expressed in monocytes and may therefore already have the chromatin accessibility state established. To test this, the accessibility of the response genes and moDC marker genes were examined in monocytes and in moDC. The monocyte marker CD14 was accessible in monocytes and closed in moDC (Figure 4B), whereas the DC marker DC-SIGN as well as HLA-DR, PD-L1 and TNFα exhibited a higher accessibility in moDC compared to the more closed configuration in monocytes (Figure 4C). The promoters of IL-6, IL-1β, and MMR had similar open accessibility in monocytes and moDC (Figure 4C). Chromatin changes are also formed by histone modifications and histone H3K4me3 and histone H3K27Ac have been associated with an active chromatin state coupled to the establishment of a trained or tolerised immune phenotype (17). HLA-DR and PD-L1 exhibited a trend of higher H3K27Ac at the promoter in cells stimulated by LPS alone (Figure 4D, upper panel), whereas no change was observed H3K4me3 (Figure 4D, lower panel). No such trends in the histone modifications were observed in other groups of gene (Supplementary Figures S4C and S4D). This suggests that no trained phenotype was induced at this short time-span and the H3K27Ac at the promoter of HLA-DR and PD-L1 may be associated with the transcriptional activation by LPS. Taken together, the genes important for a DC response already have a hyper-accessibility chromatin structure, established in differentiation steps prior to the activation of the genes upon stimulation.

**Figure 4.**
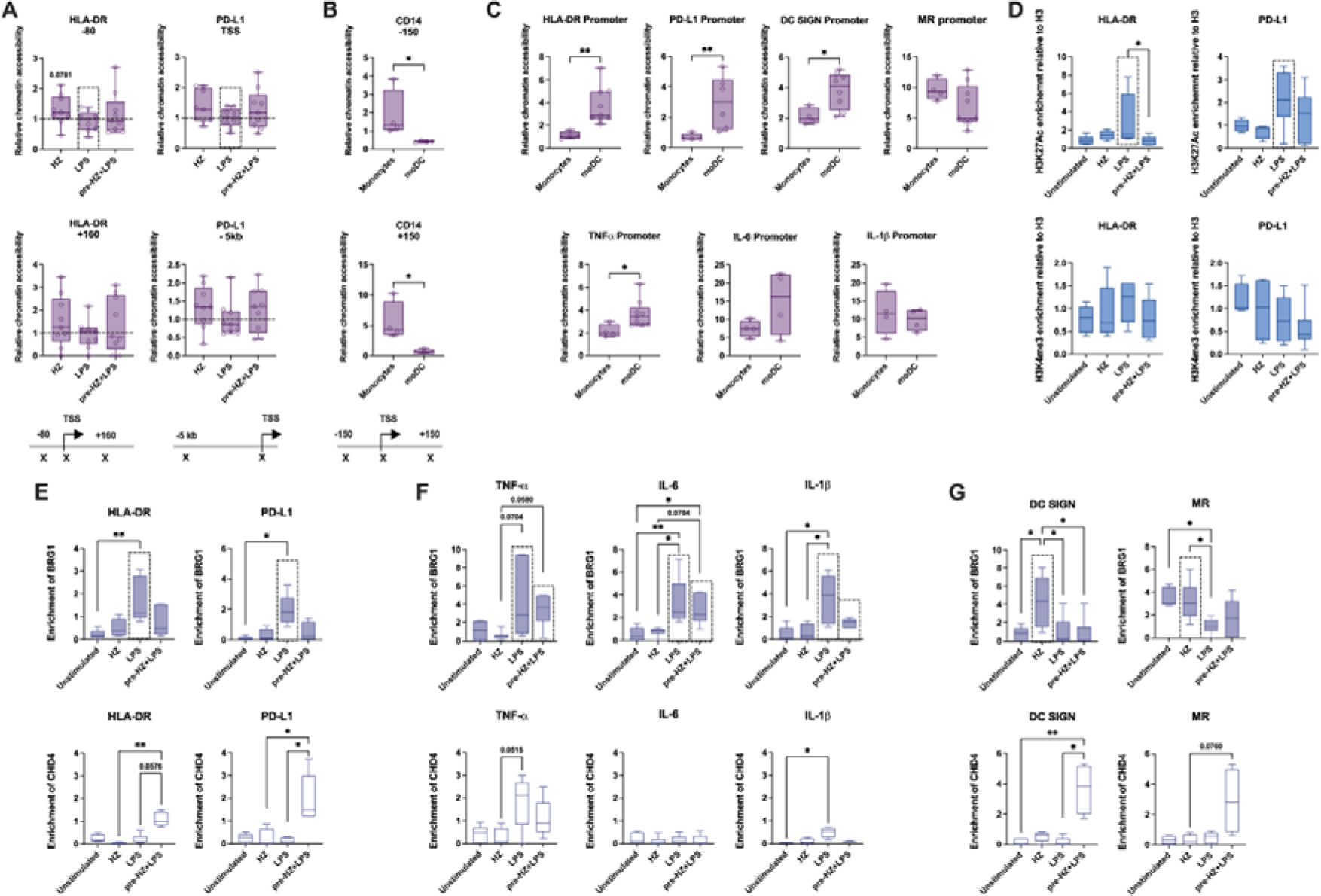
Exposure to HZ does not alter chromatin state but it changes the recruitment of BRG1 (SWI/SNF complexes) and CHD4 (NuRD) to the promoters of moDC genes. **(A)** Relative chromatin accessibility at the promoters of HLA-DR and PD-L1 genes. The primer positions are depicted under the graphs for each gene. Non-parametric Wilcoxon matched-pairs signed rank test was applied to determine significant differences, n.s.=p>0.05. Relative chromatin accessibility at the promoters of **(B)** CD14 and **(C)** HLA-DR, PD-L1, DC SIGN, MMR, TNF-α, IL-6 and IL-1β, in isolated monocytes and differentiated moDC on day 5. Non-parametric Mann-Whitney test was used to determine the differences, *p<0.05, **p<0.01. **(D)** The recruitment of histone H3K27Ac (upper graphs) and H3K4me3 (lower graphs), at the promoters of HLA-DR and PD-L1 genes. H3K27Ac and H3K4me3 were normalised to the enrichment of H3. The enrichment of BRG1 and CHD4 at the promoters of **(E)** HLA-DR and PD-L1, **(F)** TNF-α, IL-6 and IL-1β, **(G)** DC SIGN and MMR in moDC. (D-G) Non-parametric Kruskal-Wallis ANOVA test, followed by Dunńs multiple was comparison was applied to determine significant differences, n.s.=p>0.05, *p<0.05, **p<0.01. N=3-9.

### Exposure to HZ changes the recruitment of the SWI/SNF complex

Next, we examined the association of chromatin remodelling complexes to the immune genes; SWI/SNF complexes are required for transcriptional activation of many immune genes and NuRD or EZH2 in PCR2-polycomb for the repression (26, 33, 34). The ATPase BRG1/SMARCA4 was recruited to the promoters of HLA-DR and PD-L1 in LPS stimulated cells, while co-exposure with HZ and of LPS had reduced levels of BRG1 associated with the promoters, which is in agreement with the reduced levels of transcript (Figure 4E, upper panel). Co-exposure of HZ and LPS led instead to recruitment of the ATPase CHD4 in NuRD, but not the methyltransferase EZH2, to the promoters (Figure 4E, lower panel and Supplementary S4E). The promoters of the inflammatory genes, TNFα, IL-6, IL-1β, and CCL5, displayed higher BRG1 occupancy in cells stimulated with LPS and most genes retained this high level in cells co-exposed with HZ and LPS (Figure 4F, upper panel and Supplementary Figure S4F, upper panel). CHD4 was only associated with the TNFα promoter in LPS stimulated cells, indicating that repression of the transcription had started on early primary response genes (Figure 4F, lower panel and Supplementary Figures S4F, lower panel). The promoters of IL-23, IL-10, and CD83 did not display any changes in the association with BRG1 or CHD4 (Supplementary Figure S4G), suggesting that other factors are operating on these gene promoters. In contrast to the other groups of gene, BRG1 levels associated to the promoters of DC-SIGN and MMR were higher in cells exposed to HZ alone, which maintained a higher transcriptional level than in LPS-stimulated cells, and CHD4 associated with the promoters in cells co-exposed with HZ and LPS (Figures 4G), possible to allow for a reduction in transcription.

### SWI/SNF variants operating on the genes decide transcription factor preference

To further investigate the specificity of the binding of factors to the different types of gene, we determined the expression of ncRNAs regulating the binding of BRG1-SWI/SNF to immune genes. The recruitment of SWI/SNF complexes to immune genes is affected by two LPS-regulated lncRNAs, IL-7- as and FIRRE, in opposite ways; IL-7-as enhances, and FIRRE reduces the association (36, 37). Both RNA levels were significantly induced by LPS stimulation and co-exposure to LPS and HZ for 2 h compared to HZ exposed and unstimulated cells (Figure 5A), suggesting that the expression of these RNAs does not explain the differential recruitment of factors to the promoters. Instead, we examined whether the promoters associated with different SWI/SNF complex constellations and used BRD9 as a marker for the non-canonical SWI/SNF (ncBAF complex), BAF180/PBRM1 for PBAF and BAF250/ARID1 for the conventual BAF (cBAF) complex. BAF180 was recruited to the HLA-DR and PD-L1 promoters upon LPS stimulation for 2 h (Figure 5B), something that HZ exposure clearly prevented at the PD-L1 promoter. This demonstrates that these genes worked with the PBAF constellation to activate transcription and associated with NuRD in co-exposed cells to decrease transcription. The promoters of inflammatory genes TNFα, IL-6 and IL-1β showed a preference for BRD9, which associated upon LPS stimulation and co-exposure (Figure 5C). BAF250 was only found associated with CCL5 in cells stimulated with LPS and co-exposed cells (Figure 5D), which indicates that the cBAF complex is not involved in the HZ affected gene in the early response. HZ-exposure to DC-SIGN and MMR led to both BRD9 and BAF180 associating at their promoters (Figure 5E), both subunits were lost in the co-exposed cells together with BRG1 and replaced by CHD4. In conclusion, the specificity in the transcriptional response to HZ is achieved by the recruitment of specific SWI/SNF complexes, with different features, to the various promoter.

**Figure 5.**
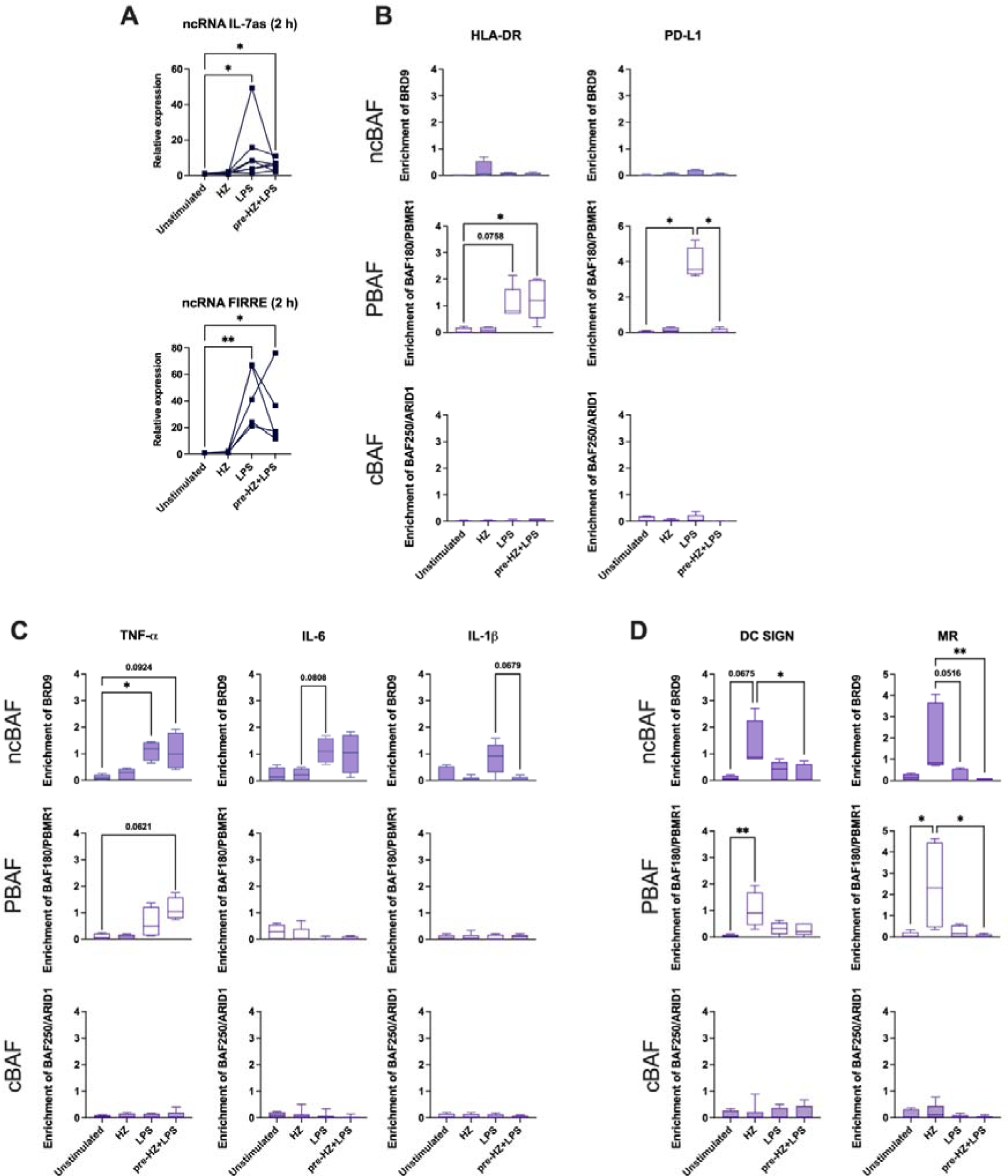
Differential recruitment of SWI/SNF complex signature proteins BRD9, BAF180 /PBMR1 and BAF250/ARID1 at the promoters of moDC genes. **(A)** Relative expression of ncRNA IL-7as and ncRNA FIRRE in moDC after 2 h exposure. The enrichment of BRD9 (ncBAF), BAF180/PBMR1 (PBAF) and BAF250/ARID1 (cBAF) proteins at the promoters of **(B)** HLA-DR and PD-L1, **(C)** TNF-α, IL-6 and IL-1β, **(D)** DC SIGN and MMR. (A-D) Non-parametric Kruskal-Wallis ANOVA test, followed by Dunńs multiple was comparison was applied to determine significant differences, n.s.=p>0.05, *p<0.05, **p<0.01, n=3-7.

## Discussion

Malaria co-infection with bacteria accelerate both the malaria progression and the bacterial infection (38, 39). The cellular response to malaria antigens and its response during co-infections are multifactorial and contradictory results have been reported (2). Here, we focused on the immune modulatory effects imposed by HZ during co-infections with bacteria and to answer the question, we investigated the early transcriptional events in DCs. The transcriptome analysis of the early gene expression response to iRBC in blood-derived DC shows that the main group of gene affected is associated with lipid synthesis and only a few immune genes were upregulated (7). Here, we show that although HZ had no direct effects on gene expression of immune genes, it modulated the response to bacterial stimuli. HZ did not induce the maturation of moDC and the cells did not secrete inflammatory cytokines and chemokines early in the acute phase after exposure. Instead, HZ exposure hampered the early transcriptional activation induced by LPS of HLA-DR, PD-L1 and CXCL10 and at the same time maintained the expression of the immaturity markers DC-SIGN and MMR, possibly to dampen or skew a subsequent T-cell response. The transcriptional response is regulated by activated transcription factors and in addition to activating NFκB signalling (21–23), we show that HZ activated IRF3 signalling after 2 h. Even if activated, the factors were differentially recruited to the different genes; activated IRF3 was differentially recruited to the promoters of DC-SIGN and MMR and not to other IRF3 targets, such as CCL5. Differential recruitment may occur in several ways, and HZ is a molecule able to modulate the binding of transcription factors by replacing activating SWI/SNF complexes by the NuRD complex during co-infection on specific genes such as HLA-DR and PD-L1 (Figure 6).

**Figure 6.**
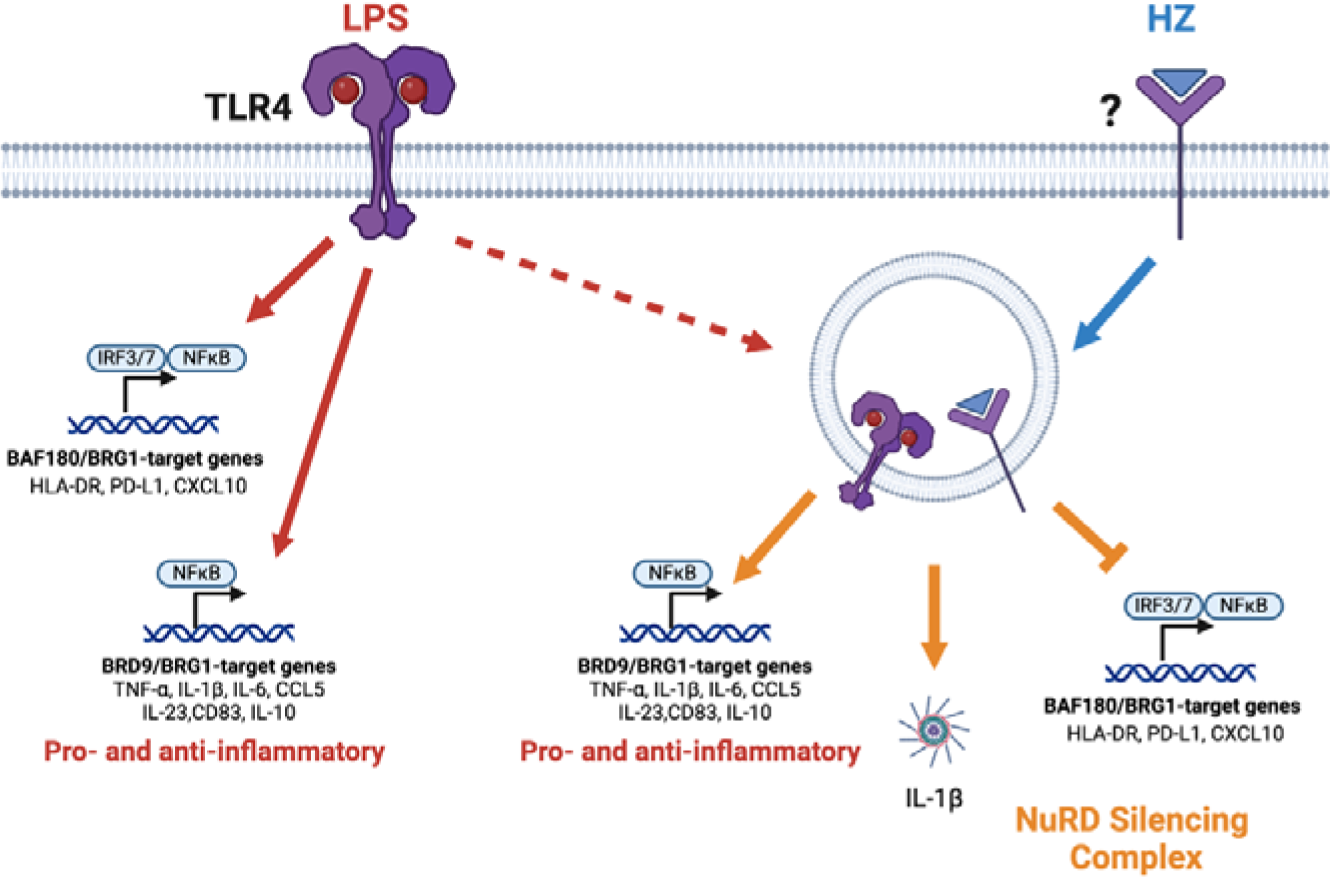
Model of the induction of LPS through TLR4 alone to the right and co-exposure with LPS and opsonized HZ leading to NuRD dampening the LPS response on HLA-DR and PD-L1 to the left.

The specific response of genes depends on their promoter and enhancer architecture as well as on their chromatin state. Immune genes are divided into groups depending on the type of response elements in regulatory regions and HLA-DR, the PD-L1 as well as the CXCL10 are interferon stimulated genes (IGS) (40–42). The promoters of these genes harbour interferon-sensitive response elements that are specifically responsive to IFNγ activation of the transcription factor STAT1 and IRFs (43, 44). The gene promoters also harbour NFκB binding sites and are regulated by TLR signalling, through the NF-κB pathway in response to pathogens. Inflammatory and maturation genes are also induced early by NFκB transcription factors RELA and later by IRF3/IRF7 (45, 46). While LPS stimulation recruited RELA to both IGS- and inflammatory genes, co-exposure with HZ reduced the recruitment of RELA to the IGS genes. This may be attributed to different signalling codons, such as stimulus specific activation of pathways, and scRNA transcription analyses have shown that although genes respond to the same transcription factors, they are activated at different time points and at different amplitudes (41, 45, 46). This is caused by specific responses to the type and quality of signal triggered by the stimulus, to the signals secreted by neighbouring cells and the chromatin states of individual cells (42, 46–48).

Only very small changes in chromatin accessibility or histone modification histone H3K4me3 and histone H3K27A were detected at the promoters of the genes upon activation compared to unstimulated cells after 2 h. Accessible chromatin and active histone modifications are often correlated with active transcription, but these can be set at different stages. Exposure of either iRBC or HZ for 24 hours induces a trained phenotype with increased enrichment of H3K4me3 at specific genes in monocytes (19. 20), which suggests that longer exposure times than 2 hours are required to establish training or tolerance. Furthermore, the differentiation also see changes in histone modifications and differentiation of monocytes to moDC involves loss of H3K4me3 at the promoter of the monocyte marker CD14 gene and a gain at the DC-SIGN gene (49). Similarly, the accessibility of gene promoter is pre-set during differentiation (50), which includes immune genes genomically primed at an earlier step in oligodendrolia development (51). We investigated the accessibility during development and show that the CD14 promoter changed from a more open configuration in monocytes to a closed one in moDC, and the DC-SIGN showed the opposite pattern and obtained a more accessible configuration. HLA-DR, PD-L1 and the inflammatory genes, which are expressed upon maturation either gained an open configuration during the differentiation to moDCs or already had an open configuration established in an even earlier step. Our results show that maturation of DCs does not require further alteration of accessibility at the promoters in the early acute phase and the transcriptional induction of DC depends on the activation and binding of transcription factor.

Many immune genes require SWI/SNF complexes for transcriptional activation. This can be counteracted in different ways, either through the NuRD complex that antagonises SWI/SNF activation (26, 52) or through EZH2-PCR2 that can silence the genes (34, 35, 52). We observed that the promoters of several of the investigated genes, including HLA-DR and PD-L1, associated with BRG1 when activated by LPS, and that BRG1 was replaced by the CHD4 when transcription was reduced following HZ and LPS co-exposure. These complexes are tightly correlated with chromatin changes, something we did not observe in our study. However, studies with perturbed SWI/SNF and NuRD factors show that the transcriptional effect is not always the expected (52–56). It has been shown that SWI/SNF and NuRD are involved in transcriptional changes by regulating RNA pol II occupancy at the gene promoter and in the gene body of highly expressed genes instead of chromatin remodelling (52). We did not detect EZH2 on HLA-DR and PD-L1 promoters, which confer Polycomb silencing in cancer cells (34, 35, 56). EZH2 has also been associated with silencing of SWI/SNF activated bivalent genes expressed at a low level by reducing the chromatin accessibility (52). In connection with our finding, we propose that many genes in the acute phase are induced by activated RELA and IRF3 and regulated by the SWI/SNF and the NuRD complexes through alteration of RNA pol II kinetics rather than changing chromatin configuration. In this context, HZ exposure establishes a different balance of the complexes at the HLA-DR and PD-L1 genes as well as the C-lectin genes which results in a compromised gene expression in response to concomitant LPS stimulation.

The specificity of the response was achieved by recruiting different SW/SNF constellations. These constellations operate on different regulatory elements and regulate cell type specific expression (57, 58). Our results show that these constellations worked with different gene promoters, and the BAF180-PBAF was associated with the promoters of HLA-DR and PD-L1 upon LPS stimulation. It was replaced by the NuRD complex in co-exposed cells resulting in reduced transcription. PBAF association to promoters often lead to repression, such as repression of genes in T-regs and IFNγ response genes (24, 25, 59–61) In contrast to the PBAF operating on HLA-DR and PD-L10, we found that a number of the inflammatory genes associated with the BRD9-ncBAF upon LPS stimulation, and this complex was not antagonised by NuRD upon co-exposure with HZ. We propose that the gene architecture determines what type of SWI/SNF complex that associate with the promoter and this determines the regulatory events. PBAF is associated with the NuRD antagonistic activity which may be responsible for the more repressive activity associated with the PBAF complex.

IL-1β was the only cytokine secreted from co-exposed cells after 24 h of stimulation and not from HZ and LPS alone. This suggest that neither HZ or LPS alone could activate the inflammasome, which requires the transcriptional activation of the IL-1β gene and other inflammasome genes, like NLPR3 and caspases, as well as the activation and assembly of the inflammasome (62). The activation of the inflammasome has been attributed to HZ by activating the STAT pathway through LYN/SYK kinases (30) or to induce the production of ROS (23, 30). The release of IL-1β after 24 h co-exposure with HZ and LPS may have implications in malaria co-infections with bacteria and contribute to an excessive inflammatory reaction. Equally important is the early modulation of the response, in which HZ-exposure of DCs maintains the cells in a partially immature state, which might lead to a skewed T-cell activation and fine-tune a homeostatic response to avoid random maturation. The modulation imposed of certain promoters by short-term HZ exposure primes the cells for subsequent or simultaneous bacterial infections and these effects may lead to a more severe disease during co-infections.

## Author Contributions

AÖF, ESE, MTB designed the study. GL, KT, AS, JQ, AO, IB performed lab work. GL, KT, AS, JQ, IB, ESE and AÖF analysed and/or finalized the data. GL performed statistical analyses. AÖF, GL and ESE wrote the paper.

