## Supplementary Figures and Table for "Malaria-derived hemozoin alters chromatin remodelling and skews dendritic cell responses to subsequent bacterial infections"

**SUPPLEMENTARY DATA**

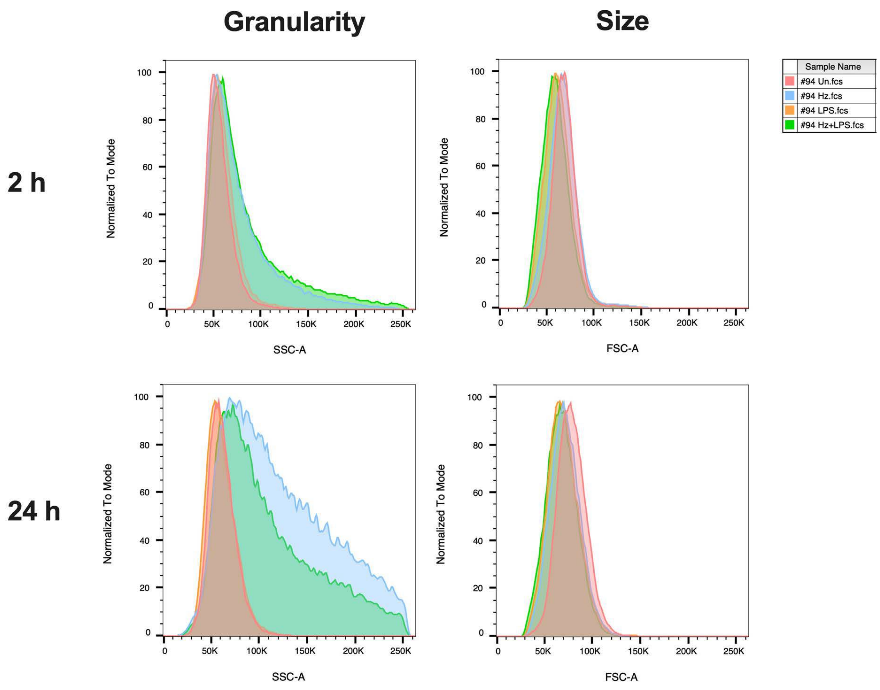


**Supplementary Figure S1. Morphology changes in moDC upon HZ exposure. (A)** Representative flow cytometry histograms showing changes in the size and granularity of moDC after they were exposed to HZ with or without LPS.

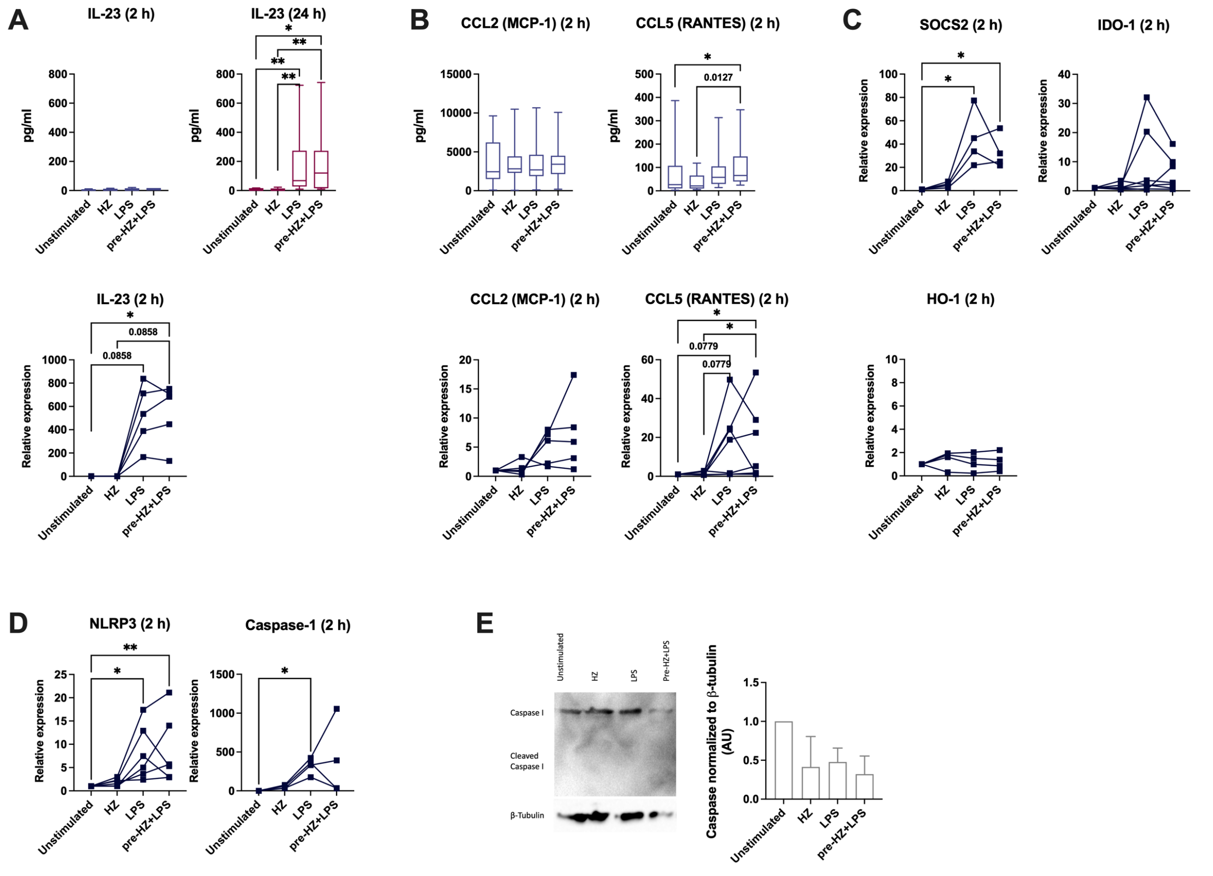


**Supplementary Figure S2. HZ does not interfere with the production of LPS-induced pro- and anti-inflammatory factors. (A)** The secretion of IL-23 from moDC after 2 h or 24 h exposure (upper panel) and relative mRNA expression of IL-23 in moDC after short term exposure (lower level), n=4-9. **(B)** The secretion of CCL2 and CCL5 (upper panel) and relative mRNA expression of CCL2 and CCL5 (lower level), in moDC after 2 h exposure, n=5-14. **(C)** Relative mRNA expression of SOCS2, IDO-1 and HO-1, n=4-7. **(D)** Relative mRNA expression levels of NLRP3 and Caspase-I, n=4-6. The mRNA levels were measured in all samples after 2 h of exposure. (A-E) Paired Friedman test followed by Dunn´s multiple comparison was used to determine statistical difference, *p<0.05, **p<0.01. **(E)** Representative immunoblot of pro-Caspase-I and cleaved Caspase-I protein in moDC after 2 h of exposure, n=3. The quantitative analysis was performed on cleaved Caspase levels normalized to β-tubulin levels (left panel).

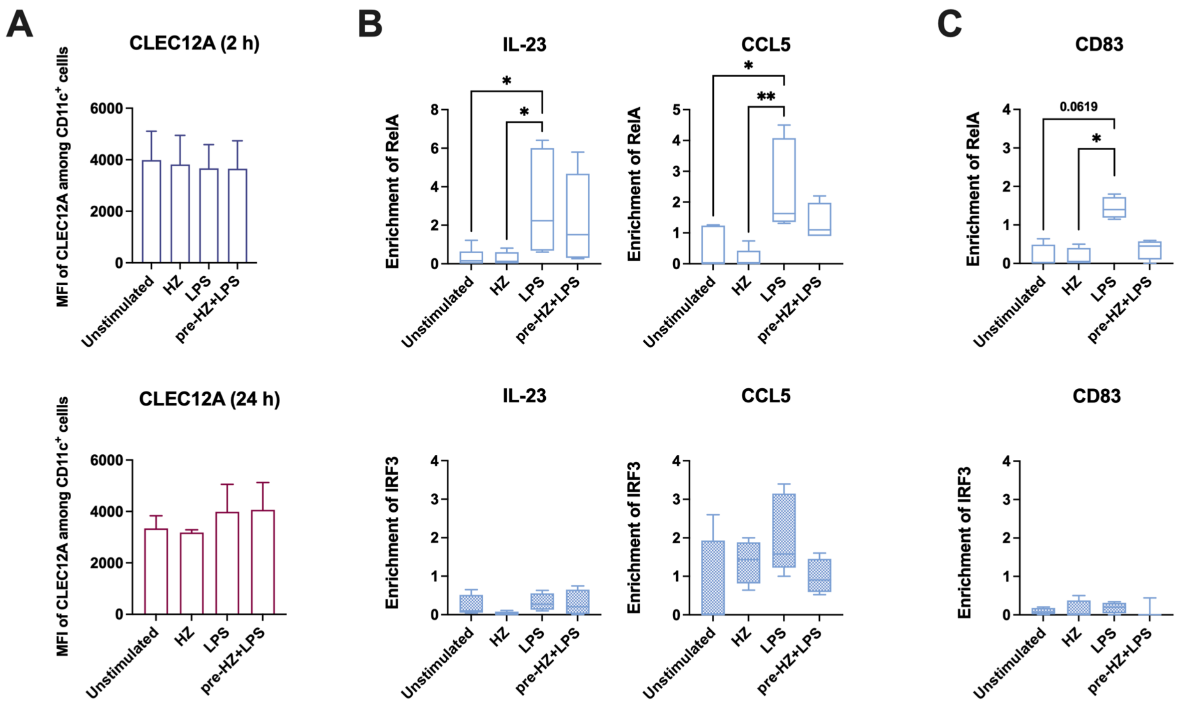


**Supplementary Figure S3. Surface expression of CLEC12A receptor and recruitment of different transcription factors at the promoters of IL-23, CCL5 genes and CD83. (A)** The mean fluorescent intensity (MFI) of CLEC12A receptor expression in moDC after 2 h exposure (upper graphs) and 24 h exposure (lower graphs), n=2-4. The data are present as median with interquartile range. The enrichment of RelA (upper panel) and IRF3 (lower panel) at the promoters of **(B)** IL-23 and CCL5 genes, n=4-7, and **(C)** CD83 gene, n=3-4. Non-parametric Kruskal-Wallis ANOVA test, followed by Dunn´s multiple was comparison was applied to determine significant differences, *p<0.05, **p<0.01.

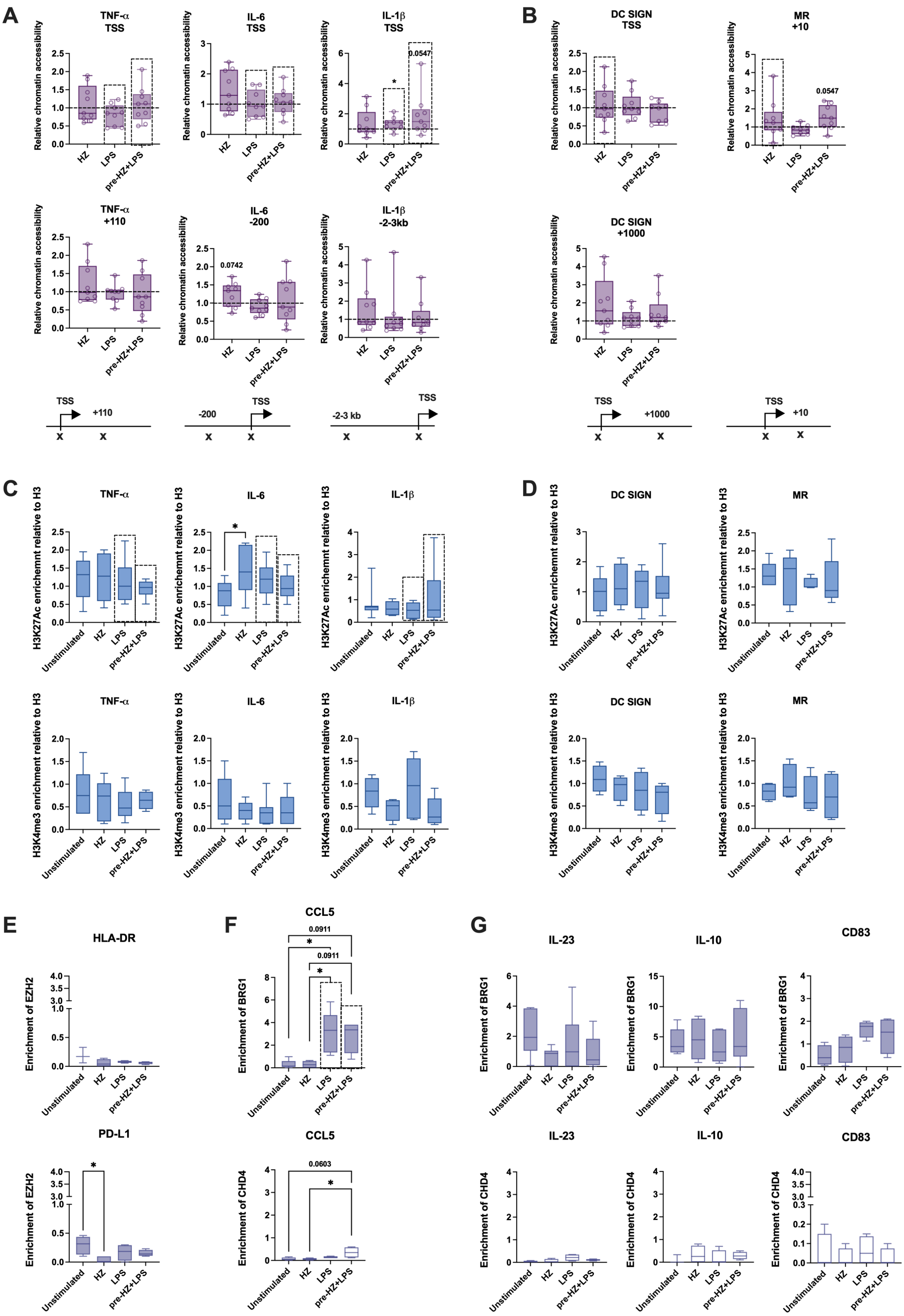
A
TNF-α TSS
IL-6 TSS
IL-1β TSS
TNF-α +110
IL-6 -200
IL-1β -2-3kb
Relative chromatin accessibility
HZ
LPS
pre-HZ+LPS
*
0.0547
0.0742
TSS
+110
-200
-2-3 kb
B
DC SIGN TSS
MR +10
DC SIGN +1000
+1000
+10
C
TNF-α
IL-6
IL-1β
H3K27Ac enrichemnt relative to H3
H3K4me3 enrichement relative to H3
Unstimulated
D
DC SIGN
MR
E
HLA-DR
PD-L1
Enrichment of EZH2
F
CCL5
Enrichment of BRG1
Enrichment of CHD4
0.0911
0.0603
G
IL-23
IL-10
CD83

**Supplementary Figure S4. Chromatin states; chromatin accessibility and the recruitment of histone modifications H3K27Ac and H3K4me3 as well as of BRG1 (SWI/SNF complexes), and CHD4 (NuRD) at the promoters of moDC genes.** Relative chromatin accessibility at the promoters of **(A)** TNF-$\alpha$, IL-6 and IL-1$\beta$ genes, n=9, and **(B)** DC SIGN and MR genes, n=9. The position of the primers is depicted under the graph. Non-parametric Wilcoxon matched-pairs signed rank test was applied to determine significant differences, n.s.=p>0.05. The recruitment of histone H3K27Ac (upper panel) and H3K4me3 (lower panel), at the promoters of **(C)** TNF-$\alpha$, IL-6 and IL-1$\beta$, n=6-11, and **(D)** DC SIGN and MR, n=4-10. **(E)** The enrichment of EZH2 at the promoters of HLA-DR and PD-L1 genes, n=3-5. The enrichment of BRG1 and CHD4 at the promoters of **(F)** CCL5, n=4-5, and **(G)** IL-23, IL-10, and CD83, n=3-9. (C-G) Non-parametric Kruskal-Wallis ANOVA test, followed by Dunn´s multiple was comparison was applied to determine significant differences, n.s.=p>0.05, *p<0.05.

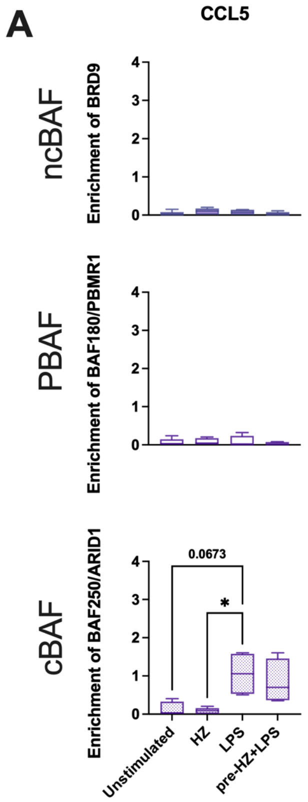


**Supplementary Figure S5. Recruitment of different SWI/SNF complex signature proteins at the promoters of genes encoding CCL5. (A)** The enrichment of BRD9, BAF180/PBMR1 and BAF250/ARID1 at the promoters of CCL5 in moDC after short term exposure. Non-parametric Kruskal-Wallis ANOVA test, followed by Dunn´s multiple was comparison was applied to determine significant differences, n.s.=$p>0.05$, *$p<0.05$, $n=4\text{-}5$.

**Table S1. Primer pair sequences.**

| Gene | Forward (F) / Reverse (R) | Oligonucleotides | References |
|---|---|---|---|
| **cDNA** | | | |
| DC SIGN | F | TTG TTG GGC TCT CCT CTG TT | [1] |
| | R | AAG TAA CCG CTT CAC CTG GA | |
| IL-6 | F | TAG AGC TTC TCT TTC GTT CCC GGT | [2] |
| | R | TGT GTC TTG CGA TGC TAA AGG ACG | |
| IL-23 | F | CTC TGC TCC CTG ATA GCC CT | [3] |
| | R | TGC GAA GGA TTT TGA AGC GG | |
| IL-10 | F | GCC TAA CAT GCT TCG AGA TC | [4] |
| | R | CTC ATG GCT TTG TAG ATG CC | |
| CD83 | F | ATT CCC TGA AGA TCC GAA AC | [5] |
| | R | GAA AAT AAC CAG AGC CAG CA | |
| CD86 | F | GTT GCC TTG AGC AAA AAC AA | [5] |
| | R | TGA GAG AGG AAG AGC TGC AA | |
| PP1A | F | AGA CAA GGT CCC AAA GAC | [6] |
| | R | ACC ACC CTG ACA CAT AAA | |
| CXCL10 | F | CCCACGTGTTGAGATCATTG | - |
| | R | TCCATCACAGCACCGGG | |
| TNF-$\alpha$ | F | TGCTTGTTCCTCAGCCTCTT | [7] |
| | R | GGTTTGCTACAACATGGGCT | |
| IL-1$\beta$ | F | TGTATGTGACTGCCCAAGATG | [8] |
| | R | TTAGTGCCGTGAGTTTCCC | |
| IL-1RA | F | TGTTAACTGCCTCCAGC | Sigma Aldrich |
| | R | ATACTTGCAAGGACCAAATG | |
| IL-18 | F | GCTGAACCAGTAGAAGACAATTGC | - |
| | R | CCAGGTTTCATCATCTTCAGCTA | |
| IL-18BP | F | GGAGGTGCTCAATGAAGGAACC | [9] |
| | R | GTGTCCAGCATTGGAAGTGACC | |
| CCR7 | F | ACAGCCTTCCTGTGTGGTTT | [10] |
| | R | ATGATGGAGTACATGATAGG | |
| CCL5 | F | AAGTCTCTAGGTTCTGAGC | Sigma Aldrich |
| | R | TTTTATGGTTGCATTGAGAAC | |
| HLA DR | F | GATTGGACCTTCCAGACCCTG | - |
| | R | ACTTGGGTGCTCCACTTGGCA | |
| MR (CD206) | F | AAATTTGAGGGCAGTGAAAG | [11] |
| | R | GGATTTGGAGTTTATCTGGTAG | |
| PD-L1 | F | TGCCGACTACAAGCGAATTACTG | [12] |
| | R | CTGCTTGTCCAGATGACTTCGG | |
| NLRP3 | F | GATCTTCGCTGCGATCAACA | [13] |
| | R | GGGATTCGAAACACGTGCAATA | |
| HO-1 | F | CCCCAACGAAAAGCACATCC | [14] |
| | R | AGACAGCTGCCACATTAGGG | |
| SOCS | F | TGCAAGGATAAGCGGACAGG | [15] |
| | R | CAGAGATGGTGCTGACGTGT | |
| IDO-1 | F | TCTGGCCAGCTTCGAGAAAG | [16] |
| | R | AGAACTAGACGTGCAAGGCG | |
| IL-7as | F | TCCTCCCCTGAATCTTCCATTAGTC | [17] |
| | R | CCTGGGCAACAGAATGTGACCTT | |
| FIRRE | F | CTGTGACCTCGCTTCACTTCT | [18] |
| | R | GTGGCAAAGAGCAGAAGATAGA | |

| | | | |
|---|---|---|---|
| BRG1 (SMARCA4) | F | GATGTCGATGATGAATATGGC | Sigma Aldrich |
| | R | ATCTGGTACTGTTTGAGGAC | |
| Caspase-1 | F | CAACTACAGAAGAGTTTGAGG | Sigma Aldrich |
| | R | AACATTATCTGGTGTGGAAG | |
| Caspase-4 | F | AAGCTCATCCGAATATGGAG | Sigma Aldrich |
| | R | ATTCTTCATGAGGACAAAGC | |
| Caspase-8 | F | CTACAGGGTCATGCTCTATC | Sigma Aldrich |
| | R | ATTTGGAGATTTCCTCTTGC | |
| CXCL9 | F | AGGTCAGCCAAAAGAAAAAG | Sigma Aldrich |
| | R | TGAAGTGGTCTCTTATGTAGTC | |
| CXCL10 | F | CCCACGTGTTGAGATCATTG | [19] |
| | R | TCCATCACAGCACCGGG | |
| IL-18 | F | GCTGAACCAGTAGAAGACAATTGC | [20] |
| | R | CCAGGTTTCATCATCTTCAGCTA | |
| **ChIP-qPCR** | | | |
| TNF-$\alpha$ promoter | F | CAGGCAGGTTCTCTTCCTCT | [21] |
| | R | GCTTTCAGTGCTCATGGTGT | |
| IL-1$\beta$ promoter | F | CACTCTTCCACTCCCTCC | [22] |
| | R | AGCCTCAAACCCTTCCTC | |
| IL-6 promoter | F | TAGCCTCAATGACGACCTAAG | [23] |
| | R | GTGGGGCTGATTGGAAACCT | |
| IL-23p19 promoter | F | GGCCTCATTCTGACGTCTTC | [24] |
| | R | CTGAAGGACCAGCCAGAGTC | |
| IL-1RA promoter | F | AGCATATGCAAAAGCCACGG | [25] |
| | R | ATGTGCAGAGCCTGTCTTGG | |
| DC SIGN promoter | F | ATCACAGGGTGGGAAATAA | [26] |
| | R | AGTCTTGGTTCCTTGGAGTC | |
| CD86 promoter | F | GCTCATCTTAACGTCATGTCTG | Qiagen |
| | R | ATTTAACCCTTTCCTTGCAGTT | |
| CD83 promoter | F | ACATTGGTGTCGAGTTGGAG | Qiagen |
| | R | GGTCTT CCTGGGGTGTCTC | |
| mTOR promoter | F | ATAAAGAGCGCTAGCCCGAA | [7] |
| | R | GGTCTTCCTGGGGTGTCTC | |
| IL-10 promoter | F | CTCCCCAGGAAATCAACT | [27] |
| | R | AAAAGCCACAATCAAGGT | |
| MR (CD206) promoter | F | CTGCCCAGCTGAAAAGAACT | [28] |
| | R | GTTTTTCCAGCCACCCTCAT | |
| PD-L1 promoter | F | CTTCGAAACTCTTCCCGGTG | [29] |
| | R | ACCTCTGCCCAAGGCAGCAA | |
| IDO-1 promoter | F | ACGGGCAACTTGGTTTCTTC | [30] |
| | R | CATGCAAGTCTGTGGTTCACT | |
| SOCS-1 promoter | F | TCCAGAAGAGAGGGAAACAG | [31] |
| | R | GGCGGCTCTCGCGCATGCTC | |
| HLA-DRA promoter | F | CAAAGGTAGGTGCTGAGGGA | [32] |
| | R | TCCATAGGTCTTTTCTCCAATGCT | |
| CCL5 promoter | F | GTTTTGAGGATACTCCTAACCACGAA | [17] |
| | R | ACACACAGCAAATGAATGACAGAGTT | |
| CXCL8 promoter | F | GCAGAGCTGTGCCTGTTGAT | [33] |
| | R | CCTTGGCAAAACTGCACCTG | |
| CXCL10 promoter | F | AGGAGCAGAGGGAAATTCCGTAAC | [34] |
| | R | AACGTGGGGCTAGTGTGCCA | |
| **ATAC-qPCR** | | | |
| ATAC Ad1_noMX | F | AATGATACGGCGACCACCGAGATCTACAC TCGTCGGCAGCGTCAGATGTG | [35] |

| | | | |
|---|---|---|---|
| DC SIGN TSS | R | AGTCTTGGTTCCTTGGAGTC | 26 |
| DC SIGN +1000 | R | TGTGTCCACAGCCAAAAG | 26 |
| TNF-α TSS | R | CTGGTCCTCTGCTGTCCTTG | 36 |
| TNF-α +110 | R | GCTTTCAGTGCTCATGGTGT | 21 |
| IL-6 TSS | R | TTCTCTTTCGTTCCCGGTGG | 37 |
| IL-6 -200 | R | GTGGGGCTGATTGGAAACCT | 23 |
| IL-1β TSS | R | TGAAAGCCATAAAAACAGCGAGG | - |
| IL-1β -2-3kb | R | AGCCTCAAACCCTTCCTC | 22 |
| HLA-DRA -80 | R | TTTCTTCTTGGGCGCTCTGT | - |
| HLA-DRA +160 | R | TCCATAGGTCTTTTCTCCAATGCT | 32 |
| PD-L1 TSS | R | GAG GAA CAA CGC TCC CTA CC | 38 |
| PD-L1 -5kb | R | ACCTCTGCCCAAGGCAGCAA | 29 |
| CD14-150 | R | TTAGGCTCCCGAGTCAACAG | Sigma Aldrich |
| CD14+150 | R | AGGTCTAGGAGGCCCCATC | Sigma Aldrich |
| MR (CD206) +10 | R | GTTTTTCCAGCCACCCTCAT | 28 |
